# Pyramidal-cell-specific hemispheric asymmetry shapes dorsoventral CA1 dynamics during rest and exploratory behavior

**DOI:** 10.64898/2026.05.15.725448

**Authors:** Chung Sub Kim, Jackson Banks, Miswakumari Lad, Seungwoo Kang

## Abstract

The hippocampus is organized along dorsal-ventral and left-right axes, but whether and how these axes interact within defined neuronal populations across behavioral states remains unresolved. Here, we combined within-animal slice electrophysiology with dual-site fiber photometry to compare dorsal and ventral CA1 activity across contralateral hemispheric configurations in mice expressing CaMKIIα-jGCaMP8s and SynI-jRCaMP1b at distinct longitudinal sites. Ventral CA1 pyramidal neurons exhibited greater intrinsic excitability and stronger AMPAR-mediated synaptic responses than dorsal CA1 neurons. *In vivo*, CaMKIIα-defined pyramidal recordings during home cage rest revealed a left-biased event-rate asymmetry within dorsal but not ventral CA1, with no comparable asymmetry in pan-neuronal SynI recordings. Apparent dorsal-ventral differences in spontaneous event rate were therefore configuration-dependent and resolved into a hemispheric, cell-type-specific effect restricted to the CaMKIIα-defined population. Lead-lag analysis showed that dorsal-ventral temporal coordination was likewise reorganized across configurations and was restricted to pyramidal-cell-biased recordings. During open-field center entries, dorsal CA1 was preferentially recruited before entry across both configurations, whereas non-coordinated entries revealed a relative post-entry suppression of contralateral ventral CA1. Together, these findings suggest that dorsal-ventral CA1 organization cannot be inferred from hemisphere-pooled designs and identify a pyramidal-cell-specific left dorsal CA1 asymmetry as a structural feature that shapes both spontaneous activity and behaviorally driven recruitment along the longitudinal hippocampal axis.

**Significance Statement:** The hippocampus is widely understood to differ along its long axis, with dorsal regions supporting spatial processing and ventral regions supporting emotional behavior. Whether this organization interacts with the left-right axis between hemispheres has remained essentially untested, because most studies pool hemispheres or record unilaterally. Using bilateral fiber photometry in mice, we show that spontaneous activity in dorsal CA1 is left-biased and that this asymmetry is specific to excitatory pyramidal neurons. The asymmetry explains apparent dorsal-ventral differences that appear configuration-dependent under conventional analysis, and it reshapes how dorsal and ventral CA1 are recruited during open-field exploration. These findings reframe hemispheric configuration from a methodological detail into an organizational variable that should be considered when interpreting hippocampal long-axis function.

## Introduction

The hippocampus is organized along multiple functional axes, with the dorsal-ventral and left-right axes representing two of its most prominent organizational principles (Shinohara et al., 2008; Fanselow and Dong, 2010; Shipton et al., 2014; Strange et al., 2014). Along the long axis, dorsal CA1 is most often discussed in the context of spatial navigation and contextual memory, while ventral CA1 connects more heavily to limbic and hypothalamic regions and is implicated in anxiety- and motivation-related behavior (Jung et al., 1994; Moser et al., 1995; Fanselow and Dong, 2010). However, a growing body of evidence indicates that the long axis is graded rather than binary, with dorsal, intermediate, and ventral domains differing in connectivity, gene expression, and cellular identity while remaining part of an integrated network (Cembrowski et al., 2016; Cembrowski and Spruston, 2019). Whether this longitudinal organization interacts with the left-right axis at the level of *in vivo* population activity has remained largely unaddressed. Functionally, ventral CA1 neurons are preferentially recruited in anxiogenic environments (Jimenez et al., 2018), and contain behaviorally specialized ensembles for anxiety-, fear-, and salience-related processing (Hong and Kaang, 2022), whereas dorsal CA1 neurons encode stimulus value and spatial features during classical conditioning (Yun et al., 2023). Most of this work, however, has examined longitudinal organization without treating hemispheric configuration as an explicit variable. This gap matters because, if hemispheric identity modifies dorsal-ventral CA1 activity, analyses that do not account for left-right configuration may conflate longitudinal differences with hemisphere-specific recruitment. Several lines of evidence suggest that hemispheric identity is not neutral. CA3-CA1 synapses show left-right asymmetries in spine structure, postsynaptic density organization, and glutamate receptor subunit allocation that depend on the laterality of presynaptic CA3 input (Kawakami et al., 2003; Shinohara et al., 2008). These asymmetries have functional consequences: hemisphere-specific optogenetic stimulation revealed differential plasticity at left versus right CA3-CA1 inputs (Kohl et al., 2011), and long-term spatial memory depends preferentially on left CA3-CA1 transmission (Shipton et al., 2014). Whether this lateralization shapes CA1 population dynamics *in vivo* has not been directly tested. A related question is whether dorsal-ventral and hemispheric effects are expressed uniformly across CA1 populations. Calcium signals under the SynI promoter integrate activity across excitatory and inhibitory neurons (Nathanson et al., 2009; Watakabe et al., 2015), whereas signals under the CaMKIIα promoter are biased toward excitatory pyramidal neurons (Benson et al., 1992; Wang et al., 2013). Comparing these promoter-defined signals within the same animals provides a direct test of whether dorsal-ventral and hemispheric effects are preferentially expressed in pyramidal ensembles or reflect broader network activity. These questions cannot be fully resolved from *ex vivo* excitability alone, because synaptic input, inhibitory recruitment, and behavioral state reshape population output in ways not captured in slice preparations (Buzsaki and Mizuseki, 2014; McGinley et al., 2015). Behavioral context also interacts with these axes. Center entry in the open field is an approach–avoidance transition that engages ventral hippocampal output to medial prefrontal cortex and basolateral amygdala (Adhikari et al., 2010, 2011; Padilla-Coreano et al., 2016) and dorsal hippocampal spatial-processing circuits (O’Keefe, 1978; Moser et al., 1995). Whether dorsal and ventral CA1 are recruited together around this transition, whether one region leads, and whether such recruitment depends on hemispheric configuration and population identity requires direct *in vivo* measurement. Here, we tested whether dorsoventral CA1 activity is jointly shaped by hemispheric configuration, population identity, and behavioral state. We first established within the same animals that dorsal and ventral CA1 pyramidal neurons differ in intrinsic excitability and Schaffer collateral-evoked synaptic responses. We then used bilateral dual-site, dual-color fiber photometry to record dorsal and ventral CA1 activity across two contralateral configurations (LD↔RV and RD↔LV) using CaMKIIα-biased and SynI-driven indicators in the same animals. We characterized spontaneous event rates and hemispheric asymmetry during home cage rest, dorsal-ventral temporal coordination, and event-aligned calcium responses around coordinated and non-coordinated open-field center entries.

## MATERIALS AND METHODS

### Animals

Male wild-type C57BL/6J mice (Jackson Laboratory, Bar Harbor, ME, USA) were used for all experiments. Mice were 6-7 weeks old at the time of viral injection for fiber photometry experiments. Mice were housed 3-5 per cage under a 12-h light/dark cycle (lights on at 07:00, lights off at 19:00) with ad libitum access to food and water. Only male mice were used in the present study to align with prior *ex vivo* electrophysiological characterizations of dorsal-ventral CA1 in male rodents and to minimize estrous cycle-related variance in calcium signal characteristics; sex effects on the present findings remain to be determined. All procedures involving animals were approved by the Institutional Animal Care and Use Committee of Augusta University.

### Drugs

D-AP5 (Cat #0106), DNQX (Cat #0189), CGP 55845 (Cat #1248), and Gabazine (SR 95531; Cat #1262) were obtained from Tocris Bioscience (MN, USA). Stock solutions were prepared in distilled water (D-AP5, Gabazine) or dimethyl sulfoxide (DMSO; DNQX, CGP 55845) and stored at -20 °C. Final working solutions were prepared by diluting stock solutions into aCSF immediately before use; final DMSO concentrations did not exceed 0.1%. Working concentrations were 25 µM for D-AP5, 20 µM for DNQX, 5 µM for CGP 55845, and 2 µM for Gabazine.

### Viruses

The viral constructs AAV-CaMKIIα-jGCaMP8s (Addgene viral prep #176752-AAV9; 100 µL; titer ≥ 1 × 10î13 vg/mL), AAV-SynI-jRCaMP1b (Addgene viral prep #100851-AAV9; 100 µL; titer ≥ 1 × 10î13 vg/mL), and AAV-Syn-ChrimsonR-tdR (Addgene viral prep #59171-AAV9; 100 µL; titer ≥ 1 × 10î13 vg/mL) were purchased from Addgene (Watertown, MA, USA). The ChrimsonR construct was used only for post hoc validation of optogenetic stimulation parameters (Supplementary Figure 2).

### Stereotaxic microinjection and optical fiber implantation

Stereotaxic surgery was performed as described previously(Kim et al., 2025). Carprofen (5 mg/kg, s.c.) was administered 10 min before surgery. Six-week-old mice were anesthetized with isoflurane (2% in 0.5 L/min oxygen) throughout the procedure. For dual-site calcium recordings, mice received contralateral viral injections targeting dorsal CA1 (dCA1) and ventral CA1 (vCA1). For the RdCA1–LvCA1 configuration, AAV-CaMKIIα-jGCaMP8s and AAV-SynI-jRCaMP1b were injected into the right dCA1 and left vCA1, respectively; for the LdCA1–RvCA1 configuration, the injection sides were reversed. For optogenetic validation experiments (Supplementary Figure 2), a separate cohort of mice received unilateral viral injections into dCA1 with AAV-CaMKIIα-jGCaMP8s together with AAV-Syn-ChrimsonR-tdTomato to enable optogenetically evoked calcium responses. Negative control animals underwent identical optical fiber cannula implantation without viral injection. Viral infusions were performed using a Hamilton syringe fitted with a 26-gauge AS conical needle. Stereotaxic coordinates (mm from bregma) were: dCA1, AP −1.8, ML ±1.6, DV −1.4 from dura; vCA1, AP −3.2, ML ±3.4, DV −4.0 from dura. A volume of 0.4 µL per site was infused at a rate of 0.1 µL/min, and the needle was left in place for 10 min after infusion before slow withdrawal. Animals were monitored for recovery of body weight following surgery. Two weeks after viral injection, optical fiber cannulae (200 µm core diameter, 0.39 NA; RWD Life Science, Shenzhen, China) were implanted to target the same CA1 regions, with fiber tips positioned approximately 0.1-0.2 mm above the viral injection sites. For optogenetic validation experiments, a single fiber was implanted above dCA1. Cannulae were secured to the skull using stainless steel anchor screws and dental acrylic cement. Fiber photometry recordings were initiated no earlier than 2 weeks after fiber implantation (approximately 4 weeks after viral injection) to allow adequate viral expression and tissue recovery.

### Behavioral experiment

All behavioral experiments were performed under blind conditions between 16:00 and 18:00 under dim light (6.6 lux). Mice were habituated to handling before experimental recordings. Behavioral data were analyzed using RWD analysis software.

### Home cage recording

Mice were transferred from the housing room to the recording room in their home cage and allowed to acclimate for 10 min before recording. The home cage was placed under dim light (6.6 lux), and the photometry tether was connected to the implanted optical fibers. Calcium activity was then continuously recorded for 10 min while the mouse was free to move within its home cage. No experimenter intervention or external stimuli were introduced during the recording. Animals showing visible signs of distress or excessive grooming during the recording session were excluded from analysis.

### Open field test

Locomotor activity and anxiety-related exploration were assessed in an open field arena (18 × 18 × 12 inch) as previously described (Kim et al., 2025). Each mouse was placed in a center of the arena and allowed to explore freely for 6 min under dim light (6.6 lux). The center zone was defined as the central 6 x 6-inch region. A line crossing was counted when the mouse’s body center crossing a demarcation line. The arena was cleaned with 70% ethanol between animals. Animal position was tracked offline using a customized software.

### Fiber photometry recordings and analysis

Fiber photometry was performed using a commercial triple-color system (RWD Life Science, Shenzhen, China). Excitation light at 410, 470, and 560 nm was delivered through a fluorescence cube and optical patch cords to the implanted cannulae. The 410-nm channel served as an isosbestic reference for correction of motion-related and other calcium-independent fluorescence changes. The 470-nm channel captured jGCaMP8s signals, and the 560-nm channel captured jRCaMP1b signals. Light intensities at the fiber tip were maintained at approximately 5 µW for each channel to minimize photobleaching, and fluorescence was acquired at 60 Hz. Home cage recordings were obtained for 10 min, with the initial 60 s of each recording excluded from analysis to minimize transient signal instability at acquisition onset. Open field recordings were collected during the 6-min behavioral session, with the initial 30 s of each recording excluded from analysis to minimize transient signal instability at acquisition onset. Raw fluorescence signals were exported and processed offline using custom analysis scripts. Traces were smoothed using a smoothing coefficient (window, W) of 12. For each recording, the signal channel was analyzed against the simultaneously acquired 410-nm reference channel over the full recording window. Baseline correction was performed using a downsampled polynomial least-squares (PLS) fitting method with a correction coefficient of β = 8. Motion correction was then applied using the 410-nm reference channel, and corrected traces were converted to ΔF/F according to the formula (F-F₀)/F₀, where F is the motion-corrected fluorescence signal and F₀ is the baseline fluorescence estimated from the downsampled PLS fit. Corrected traces were subsequently converted to z-scores for visualization and inter-animal comparison. Calcium events were detected independently in each recording channel using a robust median absolute deviation (MAD)-based peak detection algorithm with a threshold of 3× MAD and a minimum peak distance (MPD) of 1.5 s. The MPD was selected empirically based on the decay kinetics and overall temporal profile of optogenetically evoked jGCaMP8s responses recorded under the same acquisition and preprocessing conditions, together with visual inspection of spontaneous recordings, to reduce double-counting of local maxima within the decay phase of individual calcium transients while preserving sensitivity to temporally distinct events. For each recording, peak event number, event frequency, peak mean amplitude, and MAD were quantified. To assess temporal coordination between dorsal and ventral CA1, we performed a mutually closest event analysis. Peaks were detected independently in each region, and for each event in one region, the nearest event in the other region was identified based on absolute temporal proximity. Event pairs were retained only when the relationship was reciprocal, that is, each event was the nearest neighbor of the other. Matched pairs were classified as synchronous (|Δt| ≤ 1 s), dorsal-leading or ventral-leading (1-2 s window), or solo (>±2 s). These operational windows were chosen to separate near-coincident events from temporally offset events, accounting for the limited temporal precision of bulk calcium signals measured with fiber photometry; accordingly, lead-lag categories reflect temporal precedence rather than direct causal directionality(Ali and Kwan, 2020; Simpson et al., 2024).

### Optogenetic stimulation

Optogenetic stimulation was delivered using a 635-nm laser (RWD Life Science) coupled to the implanted optical fiber. Light power was measured using a power meter (Thorlabs PM100D) and adjusted to ∼1 mW at the fiber tip, accounting for transmission losses across optical components. This power level was sufficient to reliably evoke stimulus-locked calcium responses in ChrimsonR-expressing CA1 pyramidal neurons (Supplementary Figure 2). Stimulation was applied at 20 Hz (10 ms pulse width, 1 s train) for validation of optogenetic responsiveness.

### Post hoc validation of calcium indicator expression and fiber targeting

After completion of *in vivo* recordings, acute hippocampal slices were prepared for *ex vivo* validation. Calcium indicator functionality was confirmed by bath application of 30 mM KCl, which elicited robust fluorescence increases in both jGCaMP8s and jRCaMP1b channels. Viral expression and optical fiber placement were verified relative to the targeted dCA1 and vCA1 regions. Animals lacking detectable indicator expression or showing off-target expression were excluded from further analysis.

### Acute hippocampal slice preparation

Mice were anesthetized with a lethal dose of isoflurane (> 5%) in a closed plastic container and transcardially perfused with ice-cold aCSF composed of (in mM): 2.5 KCl, 1.25 NaH_2_PO_4_, 25 NaHCO_3_, 0.5 CaCl_2_, 7 MgCl_2_, 7 dextrose, 210 sucrose, 1.3 ascorbic acid, and 3 sodium pyruvate, bubbled with 95% O_2_ - 5% CO_2_. The brain was removed and hemisected along the longitudinal fissure. Dorsal hippocampal slices (300 μm, coronal plane at 10–20°) were prepared as previously described(Kim et al., 2022; Kim et al., 2025). Ventral hippocampal slices (300 μm) were prepared using a horizontal cutting angle optimized to preserve the dendritic arborization of ventral CA1 pyramidal neurons, as previously described(Kim et al., 2022). All slices were cut in ice-cold cutting aCSF using a vibrating microtome (Microslicer DTK-Zero1, DSK, Kyoto, Japan). Slices were transferred to a holding chamber containing (in mM) 125 NaCl, 2.5 KCl, 1.25 NaH_2_PO_4_, 25 NaHCO_3_, 2 CaCl_2_, 2 MgCl_2_, 12.5 dextrose, 1.3 ascorbic acid, and 3 sodium pyruvate, bubbled with 95% O_2_/5% CO_2_, and incubated at 35°C for 30 min followed by at least 45 min at room temperature before use.

### Whole-cell patch-clamp recordings

Whole-cell current-clamp recordings were performed as previously described(Kim et al., 2022; Kim et al., 2025). Briefly, hippocampal slices were submerged in a recording chamber continuously perfused with aCSF containing (in mM) 125 NaCl, 3 KCl, 1.25 NaH_2_PO_4_, 25 NaHCO_3_, 2 CaCl_2_, 1 MgCl_2_, and 12.5 dextrose, bubbled with 95% O_2_ - 5% CO_2_ at a rate of 1 ml/min and 31-33°C. CA1 pyramidal neurons were visually identified using a microscope (Olympus BX51WI, US) fitted with differential interference contrast optics(Stuart et al., 1993). Somatic patch pipettes (resistance 4–7 MΩ when filled with internal solution) were prepared with capillary glass (external diameter 1.65 mm and internal diameter 1.1 mm, World Precision Instruments) using a Flaming/Brown micropipette puller (P-1000, Sutter Instrument, CA). The internal solution contained (in mM) 120 K-gluconate, 20 KCl, 10 HEPES, 4 NaCl, 7 K_2_-phosphocreatine, 4 Mg-ATP, 0.3 Na-GTP (pH 7.3 with KOH). Recordings were obtained with a MultiClamp 700B amplifier (Molecular Devices, San Jose, CA) and acquired using pCLAMP10 software (Molecular Devices). Signals were low-pass filtered at 10 kHz, sampled at 20 kHz, and digitized via Axon Digidata 1440A (Axon Instruments). Series resistance was monitored throughout each recording; experiments were excluded if series resistance exceeded 15 MΩ. The resting membrane potential was measured as the steady-state somatic potential in the absence of injected current. Liquid junction potential was estimated to be approximately −13 mV using the Patcher’s Power Tools plug-in in Igor Pro and was not corrected. Synaptically evoked EPSPs were recorded from dorsal and ventral CA1 pyramidal neurons held at −65 mV by constant current injection. EPSP slope was measured from the initial 1 ms of the rising phase. Evoked EPSPs were elicited by bipolar tungsten electrodes placed in the stratum radiatum (SR) in the presence of D-AP5 (25 μM), gabazine (2 μM), and CGP 55845 (5 μM) to isolate AMPA receptor-mediated responses. Because dorsal and ventral CA1 pyramidal neurons differ in average apical dendritic length, with ventral neurons exhibiting longer apical dendrites than dorsal neurons(Malik et al., 2016), SR geometry is not directly equivalent along the longitudinal axis. Accordingly, the stimulating electrode was positioned in SR at approximately 150 μm from the soma in dorsal CA1 and approximately 200 μm from the soma in ventral CA1 to target a comparable portion of the apical dendritic field. Stimuli consisted of 0.1-ms current pulses at 2-8% of the maximum stimulator output (1 mA), corresponding to stimulus intensities of approximately 20-80 μA. At the end of each experiment, evoked EPSPs were confirmed to be AMPA receptor-mediated by bath application of DNQX (20 μM), which abolished the response.

### Data analysis

Input resistance was determined from the slope of the linear fit of the voltage-current (V-I) relationship between +30 and -150 pA current injections. Action potential firing in response to depolarizing current steps was quantified as the number of APs per current injection step and fitted using a linear input-output relationship. Electrophysiological data were analyzed using Easy Electrophysiology (Easy Electrophysiology Ltd., UK) and Axograph (Axograph Scientific).

### Statistical Analysis

Statistical comparisons were performed using GraphPad Prism (GraphPad Software, San Diego, CA). Normality was assessed using the Shapiro-Wilk test. Normally distributed data were analyzed by paired t-test for within-animal comparisons (e.g., dorsal vs ventral within each configuration), unpaired Welch’s t-test for between-cohort comparisons (e.g., LD-RV vs RD-LV configurations), or one-way or two-way ANOVA followed by Bonferroni post-hoc test where appropriate. Non-normally distributed data were analyzed using the Mann–Whitney U test or Wilcoxon signed-rank test. All data are presented as mean ± SEM unless otherwise indicated. Statistical significance was defined as P < 0.05. For electrophysiology experiments, n refers to the number of neurons recorded, with the number of animals indicated in the figure legends. For fiber photometry and behavioral experiments, n refers to the number of animals.

## Results

### Within-animal whole-cell recordings reveal greater intrinsic excitability and AMPAR-mediated synaptic responsiveness in ventral than dorsal CA1

The dorsal and ventral hippocampus are widely treated as functionally distinct, with dorsal CA1 implicated in spatial and mnemonic processing and ventral CA1 in emotional and motivational computations(Moser and Moser, 1998; Bannerman et al., 2004; Fanselow and Dong, 2010; Strange et al., 2014). It has been reported that ventral CA1 pyramidal neurons are intrinsically more excitable than dorsal CA1 neurons in rat(Dougherty et al., 2012; Kim and Johnston, 2015), but most of these comparisons were made between recordings from separate animals or independent slice preparations. We therefore performed whole-cell current-clamp recordings from dorsal and ventral CA1 pyramidal neurons in slices prepared from the same animals to determine how robustly this gradient is expressed within a common biological context. Voltage responses to hyperpolarizing current steps revealed clear differences in subthreshold membrane properties between dorsal and ventral CA1 (Figure 1A). Ventral CA1 pyramidal neurons showed a significantly more depolarized resting membrane potential (Figure 1B) and significantly higher input resistance (Figure 1C) than dorsal CA1 neurons. Consistent with this, depolarizing current steps elicited significantly more action potentials in ventral than dorsal CA1 neurons, and this separation persisted across the full F-I relationship, with the strongest divergence at intermediate-to-high current intensities (Figure 1D-E). These results indicate that, when measured within the same animals and under matched recording conditions, ventral CA1 pyramidal neurons are intrinsically more excitable than their dorsal counterparts. To determine whether this gradient extended to evoked excitatory synaptic drive, we recorded α-amino-3-hydroxy-5-methyl-4-isoxazolepropionic acid (AMPA) receptor-mediated excitatory postsynaptic potentials (EPSPs) in CA1 pyramidal neurons during stimulation of Schaffer collateral fibers in stratum radiatum (Figure 1F-G). Ventral CA1 neurons displayed significantly steeper EPSP slopes than dorsal CA1 neurons across stimulation intensities, with separation becoming significant at higher stimulus strengths (Figure 1H). Bath application of 6,7-dinitroquinoxaline-2,3-dione (DNQX) abolished the evoked responses (Figure 1F-G insets), confirming that the measured EPSPs were AMPAR-mediated. Together, within-animal recordings establish that ventral CA1 pyramidal neurons exhibit (1) a more depolarized resting membrane potential, (2) higher input resistance, (3) greater action potential output, and (4) stronger evoked AMPAR-mediated synaptic responses than dorsal CA1 neurons.

**Figure 1.**
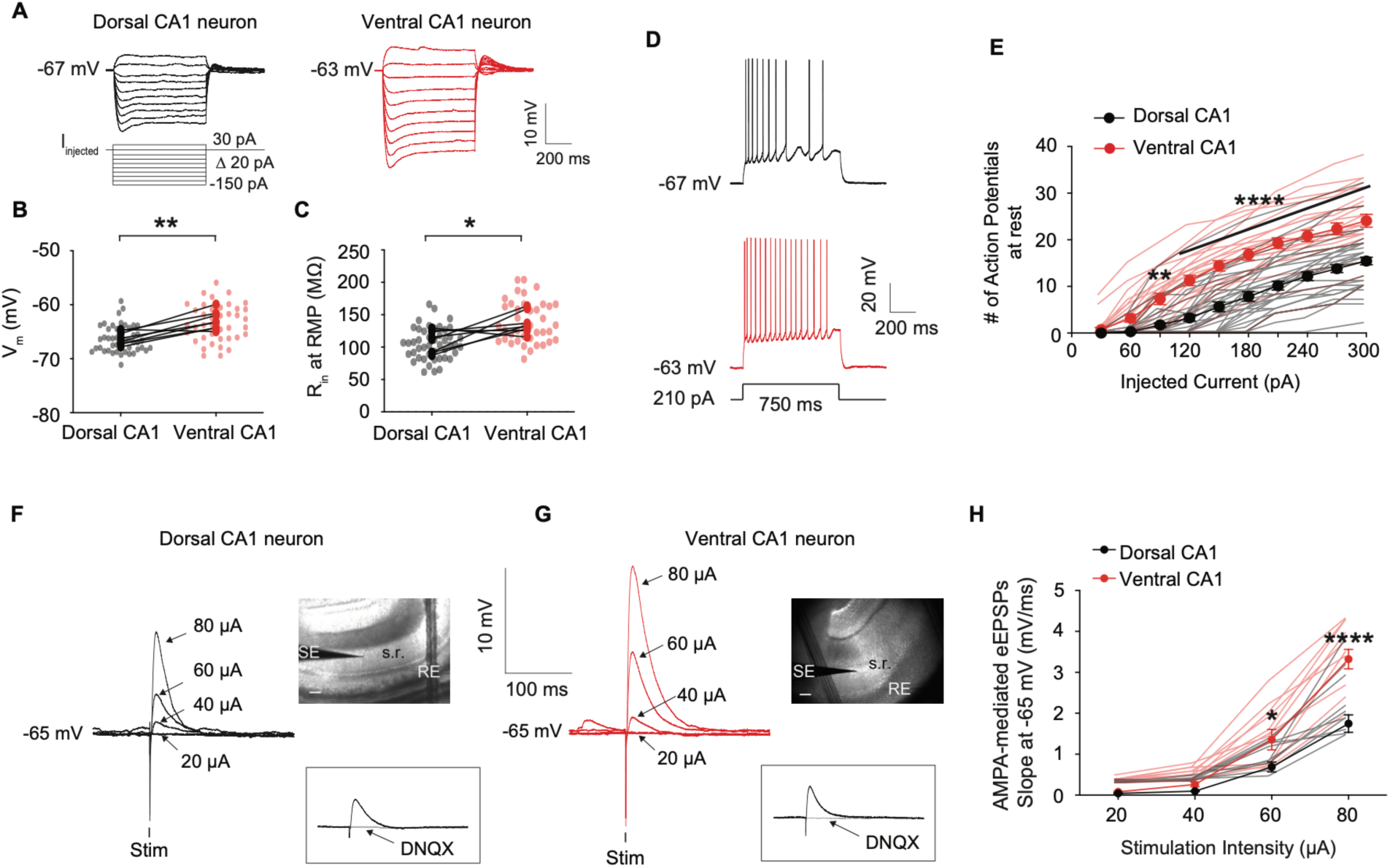
Ventral CA1 pyramidal neurons exhibit enhanced intrinsic excitability and AMPAR-mediated synaptic responsiveness compared to dorsal CA1. (A) Representative voltage responses from dorsal (black) and ventral (red) CA1 pyramidal neurons during hyperpolarizing current injections (-150 to 30 pA, Δ20 pA, 750 ms). (B) Resting membrane potential was significantly more depolarized in ventral than dorsal CA1 neurons. (C) Input resistance measured at the resting membrane potential was significantly higher in ventral CA1 neurons. (D) Representative firing patterns evoked by a 210 pA, 750 ms depolarizing current step. (E) Input-output relationship showing the number of action potentials evoked across increasing current intensities. Ventral CA1 neurons fired more action potentials than dorsal CA1 neurons across a broad range of current injections. (F, G) Representative AMPAR-mediated evoked EPSPs recorded from dorsal (F) and ventral (G) CA1 pyramidal neurons during stratum radiatum stimulation at increasing intensities (20-80 µA). Insets: AMPAR dependence confirmed by abolition of evoked EPSPs after bath application of DNQX. Scale bar: 100µm. (H) eEPSP slope quantified across stimulation intensities. Ventral CA1 neurons exhibited significantly larger evoked synaptic responses than dorsal CA1 neurons at higher stimulation intensities. Light symbols represent individual cells; bold symbols represent animal means. Lines connect dorsal and ventral animal means from the same mouse. Data are presented as mean ± SEM across animals, and all statistical analyses were performed on animal means with each animal treated as the biological replicate. Statistics: Normality was assessed using the Shapiro-Wilk test. Panels B and C: unpaired two-tailed Welch’s t-test on animal means. Panels E and H: two-way repeated-measures ANOVA (factors: region × current/stimulation intensity) followed by Sidak’s multiple-comparisons test at each intensity level. Exact p-values, test statistics, and degrees of freedom are reported in Table_statistical_analyses. *p < 0.05, **p < 0.01, ****p < 0.0001. Sample sizes: Dorsal CA1, n = 46 cells from 8 mice; ventral CA1, n = 40 cells from 8 mice.

### Validation of dual-color fiber photometry under dim home-cage conditions

Before examining how dorsal and ventral CA1 activity is organized *in vivo*, we first established that our dual-color fiber photometry system reports calcium signals reliably under the lighting conditions used throughout this study. Because RCaMP-class indicators are excited near wavelengths that overlap with ambient room light, and because home-cage exploration unavoidably exposes the recording fiber to varying illumination, we systematically characterized how arena lux affects the recorded signal (Supplementary Figure 1A-C). At 6.6 lux, the illumination used for all home cage and open field recordings reported here, fiber-tip excitation power was tuned to approximately 5 µW per channel, well within the linear regime of both jGCaMP8s(Zhang et al., 2023) and jRCaMP1b(Dana et al., 2016). Without correction, the 560 nm (RCaMP) channel showed lux-dependent baseline drift, whereas the 410 nm isosbestic reference channel remained flat (Supplementary Figure 1D). Application of PLS-based baseline correction together with isosbestic motion correction(Lerner et al., 2015) substantially attenuated these lux-dependent step shifts; however, residual lux-related drift, including a downward trend toward the end of the recording, was still detectable (Supplementary Figure 1E). For this reason, we restricted all behavioral recordings reported in this study to the lowest of the tested lux conditions (6.6 lux), at which residual artifact is smallest. We further analyzed all photometry data as Z-scored traces rather than ΔF/F, so that each recording was normalized to its own local baseline statistics, and we restricted analyses to event-rate, classification, and event-aligned AUC metrics that depend on local fluctuations around behavioral events rather than on absolute fluorescence levels across the session. All recordings were processed through the same correction pipeline. We further validated indicator-driven signal specificity using complementary negative and positive controls (Supplementary Figure 2). Animals implanted with optic fibers but injected with no virus showed no detectable response to identical optogenetic stimulation parameters (Supplementary Figure 2A-D), excluding light-evoked autofluorescence or stimulation artifact as a source of signal. In contrast, animals co-expressing AAV-CaMKIIα-jGCaMP8s and AAV-SynI-ChrimsonR-tdT(Klapoetke et al., 2014) exhibited robust, time-locked jGCaMP8s transients to 20 Hz, 10 ms-pulse, 1-second optogenetic trains (Supplementary Figure 2E-I). Together, these controls establish that the photometry recordings reported throughout this study reflect indicator-driven calcium signals.

### *In vivo* home cage activity is shaped by a CaMKIIα-specific hemispheric asymmetry within dorsal CA1, not by uniform ventral dominance

Given that ventral CA1 neurons are intrinsically more excitable than dorsal CA1 neurons *ex vivo* (Figure 1), we next examined whether this gradient was reflected in spontaneous *in vivo* activity. Because hippocampal lateralization is documented at the level of CA3–CA1 synaptic plasticity, NMDAR composition, and spatial-memory dependence(Shinohara et al., 2008; Kohl et al., 2011; Shipton et al., 2014), and because most published bilateral CA1 photometry studies pool across hemispheres, we recorded from both LD↔RV and RD↔LV configurations in the same animals (Figure 2A). Each animal received contralateral co-injections of AAV-CaMKIIα-jGCaMP8s and AAV-SynI-jRCaMP1b into dorsal and ventral CA1, allowing simultaneous comparison of pyramidal-cell-biased (CaMKIIα) and pan-neuronal (SynI) calcium signals across the same anatomical sites within the same animal, an internal cell-type-specificity control absent from most prior work in this region(Jimenez et al., 2018; Yun et al., 2023). Viral expression and indicator function were verified post hoc by KCl-evoked fluorescence in dCA1 and vCA1 slices (Figure 2B). Spontaneous CaMKIIα-jGCaMP8s activity was reliably detected at all four sites during home cage recordings (Figure 2C-D). However, the relationship between dorsal and ventral CA1 event rates depended on which hemispheres were sampled. In the LD↔RV configuration, peak event number and peak frequency did not differ significantly between dorsal and ventral CA1 (Figure 2E–F). In the RD↔LV configuration, in contrast, left vCA1 showed significantly higher event number and frequency than right dCA1 (Figure 2G–H). Taken in isolation, the latter pattern would appear consistent with an excitability-predicted vCA1 > dCA1 gradient, but the absence of the same effect in the LD↔RV configuration argues against a simple regional explanation. We therefore re-organized the data by hemisphere within each longitudinal subregion. Within dorsal CA1, left dCA1 exhibited significantly higher peak event number and peak frequency than right dCA1 (Figure 2I-J). Within ventral CA1, left and right vCA1 did not differ on either metric (Figure 2K-L). The apparent configuration-dependent dorsal-ventral difference is therefore parsimoniously explained by a hemispheric asymmetry restricted to dorsal CA1: when right dCA1 (the lower-activity site) is paired with left vCA1, vCA1 appears dominant; when left dCA1 (the higher-activity site) is paired with right vCA1, the apparent gradient collapses. Interestingly, this left-biased asymmetry was specific to the CaMKIIα-defined pyramidal-cell-biased signal. SynI-jRCaMP1b recordings(Supplementary Figure 3) showed no significant dorsal-ventral differences in either configuration (Figure 2M-P) and no left-right differences within dorsal or ventral CA1 (Figure 2Q-T). Supplementary analyses of mean peak amplitude and median absolute deviation revealed no consistent regional or hemispheric effects in either indicator (Supplementary Figure 4), indicating that the asymmetry is expressed in event rate rather than in event size or baseline signal variability. These findings constitute a triple dissociation by 1) region (dCA1 only), 2) hemisphere (left > right), and 3) cell-type marker (CaMKIIα only) and suggest two consequential conclusions. First, the *in vivo* organization of spontaneous CA1 activity does not follow directly from the *ex vivo* intrinsic-excitability gradient; if it did, vCA1 should dominate in both configurations and in both indicators, which it does not. Second, hemisphere-pooled or single-configuration designs are likely to misrepresent CA1 activity asymmetry, because the apparent dorsal-ventral relationship depends on which hemispheres are sampled.

**Figure 2.**
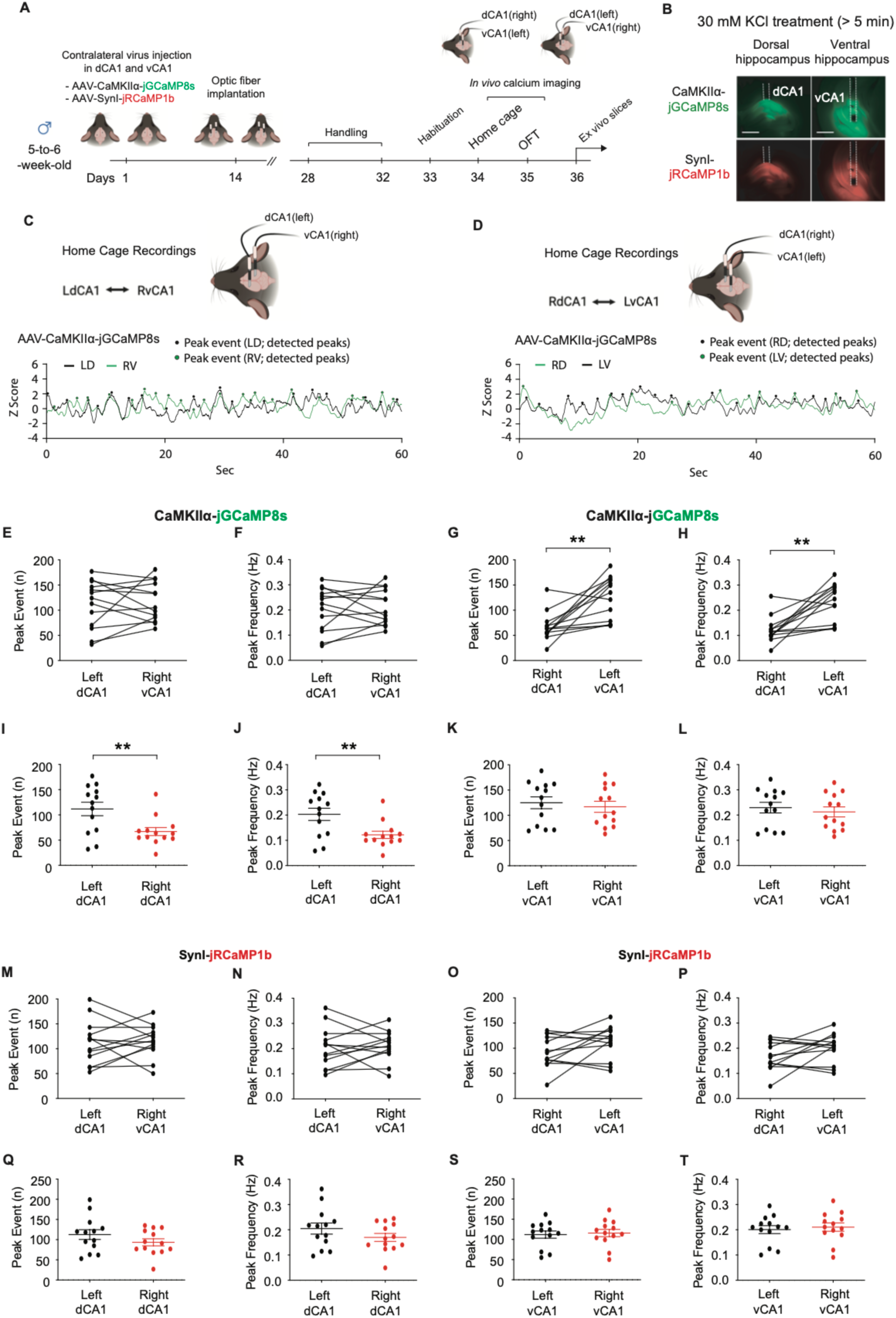
Home cage recordings reveal a CaMKIIα-specific dorsal CA1 hemispheric asymmetry and configuration-dependent dorsoventral organization. (A) Experimental timeline. Five-to-six-week-old male C57BL/6J mice received contralateral viral injections of AAV-CaMKIIα-jGCaMP8s and AAV-SynI-jRCaMP1b into dorsal CA1 (dCA1) and ventral CA1 (vCA1), followed by optic fiber implantation. Dual-site fiber photometry recordings were obtained during home cage behavior on day 35, after handling and habituation, and *ex vivo* validation was performed on day 36. (B) Representative fluorescence images of jGCaMP8s (green) and jRCaMP1b (red) expression in dorsal and ventral hippocampus. KCl stimulation (30 mM, > 5 min) evoked robust fluorescence increases at both indicator wavelengths, confirming sensor functionality at all four recording sites. Scale bar: 500µm. (C, D) Representative 60-s Z-scored CaMKIIα-jGCaMP8s calcium traces from contralateral dorsal-ventral CA1 recordings during home cage behavior in the LdCA1↔RvCA1 (C) and RdCA1↔LvCA1 (D) configurations. Detected calcium events are marked. (E, F) In the LdCA1↔RvCA1 configuration, peak event number (E) and peak frequency (F) did not differ significantly between dorsal and ventral CA1. (G, H) In the RdCA1↔LvCA1 configuration, LvCA1 showed significantly higher peak event number (G) and peak frequency (H) than RdCA1. (I, J) Hemispheric comparison within dorsal CA1. Left dCA1 exhibited significantly higher peak event number (I) and peak frequency (J) than right dCA1, indicating a CaMKIIα-defined dorsal CA1 hemispheric asymmetry. (K, L) Left and right ventral CA1 did not differ in peak event number (K) or peak frequency (L). (M-P) SynI-jRCaMP1b recordings showed no significant dorsoventral differences in either the LdCA1↔RvCA1 (M, N) or the RdCA1↔LvCA1 (O, P) configuration. (Q-T) SynI-jRCaMP1b recordings showed no significant left-right differences within dorsal CA1 (Q, R) or within ventral CA1 (S, T). Data points represent animal-level values. Connected lines in panels E-H and M-P indicate paired dorsal-ventral recordings obtained simultaneously from the same animal. Data are presented as mean ± SEM. Statistics: Normality was assessed using the Shapiro-Wilk test. Within-animal dorsoventral comparisons (E-H, M-P) were analyzed using paired two-tailed t-tests. Hemispheric comparisons (I-L, Q-T) compared signals across animals between the two configurations and were analyzed using unpaired two-tailed Welch’s t-tests. Exact p-values and test statistics are reported in Table_statistical_analyses. **p < 0.01. Sample sizes. LdCA1↔RvCA1, n = 13 mice; RdCA1↔LvCA1, n = 13 mice.

### Pyramidal-cell-biased temporal coordination between dorsal and ventral CA1 is configuration-dependent

We next asked whether the configuration-dependence observed in event rate (Figure 2) extended to the temporal relationship between dorsal and ventral CA1 events. We used an operational reciprocal nearest-neighbor classification adapted from prior multi-region calcium-event analyses(Ali and Kwan, 2020; Simpson et al., 2024), in which detected peaks in each recording channel were paired with their mutually closest event in the partner channel. Pairs were classified according to the inter-event interval Δt as: synchronous (|Δt| ≤ 1 s), dorsal-leading (vCA1 follows dCA1 by 1-2 s), ventral-leading (dCA1 follows vCA1 by 1-2 s), or solo (no reciprocal partner within ±2 s) (Figure 3A–B). These categories describe temporal precedence, not causal direction. Total event yield did not differ between configurations or between indicators (Figure 3C), ruling out detection bias as a source of category-level differences. In the CaMKIIα-jGCaMP8s recordings, the proportion of synchronous events was significantly lower in the RD↔LV configuration than in the LD↔RV configuration (Figure 3D), and ventral CA1 solo events were correspondingly higher in RD↔LV (Figure 3E). The percentages of dorsal-leading, ventral-leading, and dorsal CA1 solo events did not differ significantly between configurations (Figure 3F-H). The temporal effect was therefore not a shift toward systematically dorsal- or ventral-leading activity but rather a loss of bilateral synchrony coupled to a gain of independent ventral CA1 events when right dCA1 was the dorsal recording site. In SynI-jRCaMP1b recordings, none of these temporal categories differed significantly between configurations (Figure 3D–H). The configuration-dependent reorganization of dorsal-ventral coupling was therefore restricted to the CaMKIIα-defined pyramidal-cell-biased population, paralleling the cell-type specificity of the event-rate asymmetry in Figure 2. Sensitivity analyses using 750 ms synchrony windows yielded the same qualitative pattern (data not shown), indicating that this result is not specific to the 1-s threshold.

**Figure 3.**
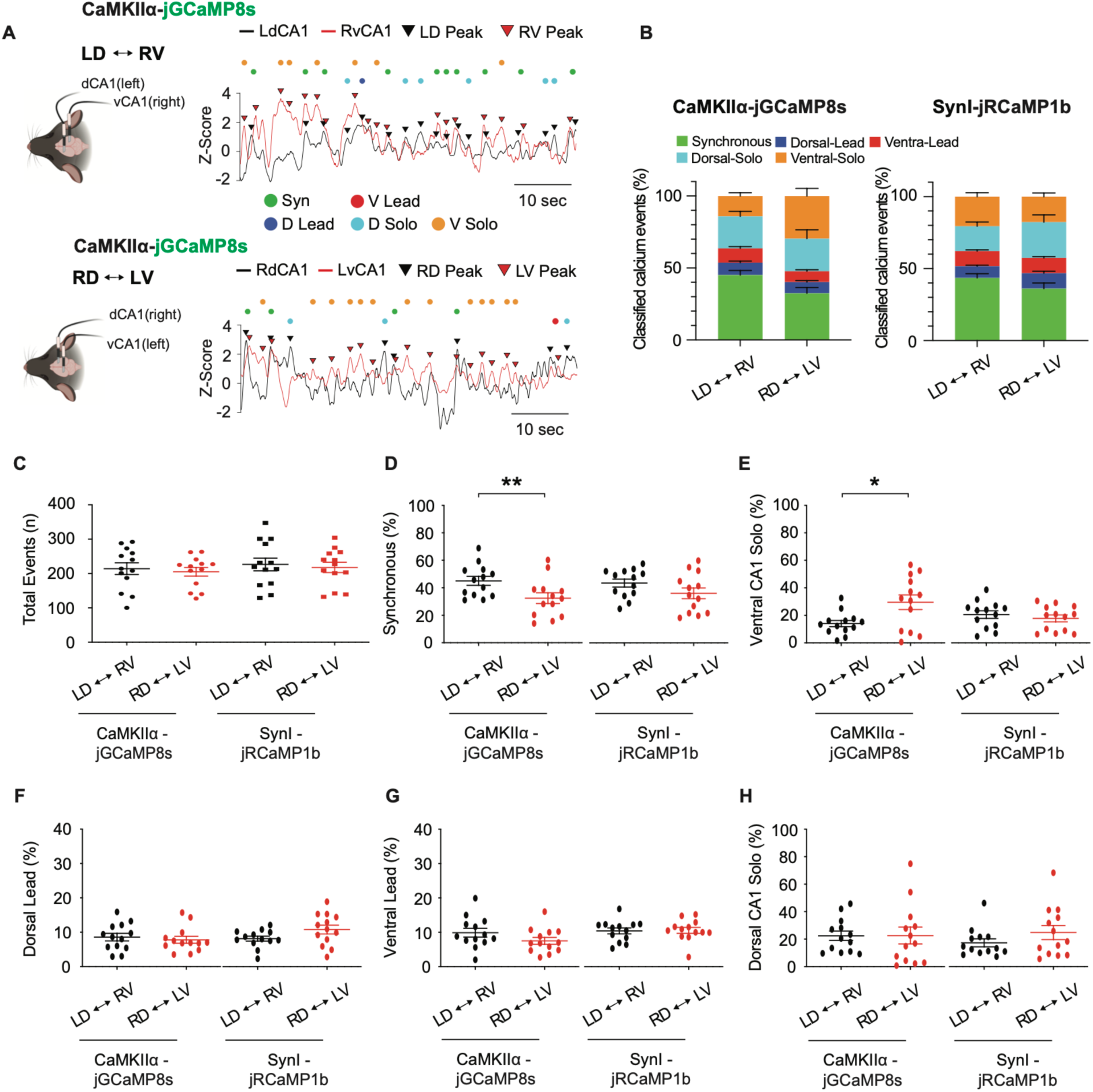
Dorsal-ventral CA1 temporal coordination during home cage behavior is configuration-dependent in the CaMKIIα-defined population. (A) Representative dual-site CaMKIIα-jGCaMP8s calcium traces from the LdCA1↔RvCA1 (top) and RdCA1↔LvCA1 (bottom) configurations during home cage behavior. Detected events were classified using a reciprocal nearest-event matching algorithm: synchronous (|Δt| ≤ 1 s), dorsal-leading (vCA1 follows dCA1 by 1-2 s), ventral-leading (dCA1 follows vCA1 by 1-2 s), or dorsal/ventral solo (no reciprocal partner within ± 2 s). These categories describe temporal precedence at the population level and do not infer single-cell timing or causal direction. (B) Stacked distribution of classified calcium events for CaMKIIα-jGCaMP8s and SynI-jRCaMP1b recordings across configurations. (C) Total number of detected calcium events did not differ across configurations or indicators, indicating that category-level differences are not driven by detection bias. (D) Synchronous events. CaMKIIα-jGCaMP8s recordings showed a significant reduction in synchronous events in the RdCA1↔LvCA1 configuration; SynI-jRCaMP1b recordings showed no significant configuration effect. (E) Ventral CA1 solo events were significantly increased in the RdCA1↔LvCA1 configuration in CaMKIIα-jGCaMP8s recordings, indicating greater independent ventral CA1 activity. (F, G) Dorsal-leading (F) and ventral-leading (G) event proportions did not differ significantly between configurations in either indicator. (H) Dorsal CA1 solo event proportions did not differ significantly between configurations in either indicator. Data points represent individual mice. Data are presented as mean ± SEM. Five non-orthogonal event categories were tested per indicator (synchronous, dorsal-leading, ventral-leading, dorsal solo, ventral solo). Exact p-values are reported in Table_statistical_analyses. *p < 0.05, **p < 0.01. Sample sizes. LdCA1↔RvCA1, n = 13 mice; RdCA1↔LvCA1, n = 13 mice.

### Coordinated open-field center entries recruit dorsal CA1 ahead of ventral CA1

Home cage recordings were obtained under unconstrained conditions in which mice cycled between rest and brief locomotor bouts. To ask whether the same dorsal-ventral organization was expressed around a defined behavioral transition, we used center entry in the open-field test as a time-locked event. Center entry is an approach-avoidance transition that engages ventral hippocampal output to mPFC and BLA (Adhikari et al., 2010, 2011; Padilla-Coreano et al., 2016) and that has been used to dissociate dorsal and ventral CA1 contributions to anxiety-related behavior(Felix-Ortiz et al., 2013; Jimenez et al., 2018). Around center entries, dorsal and ventral CA1 frequently showed coordinated calcium activity, but coordination was heterogeneous across entries (Figure 4A-B). We separated entries into synchronous events (a peak was detected in both recorded regions within ±1 s of entry onset) and nonsynchronous events (no bilateral peak within this window) and analyzed the two classes separately. Because both classes share the same behavioral definition (i.e., entry into the central zone), they constitute paired conditions for the same locomotor transition. For synchronous entries in the LD↔RV CaMKIIα-jGCaMP8s recordings, the event-aligned signal rose in both regions, but the pre-entry AUC was significantly greater in left dCA1 than in right vCA1 (Figure 4C-D). Post-entry AUC, peak amplitude, and peak time did not differ (Figure 4E-G), indicating that the dorsal bias was carried by the integrated pre-entry response rather than by a difference in peak magnitude or latency. The same pre-entry dorsal bias appeared in the RD↔LV configuration, where right dCA1 also exceeded left vCA1 in pre-entry AUC (Figure 4H-I). In RD↔LV, however, the dorsal bias additionally extended into the post-entry window (Figure 4J), with peak amplitude and peak time again unchanged (Figure 4K-L). The CaMKIIα-defined population therefore shows a consistent dorsal-leading recruitment around coordinated center entries, with the temporal extent of that bias modulated by recording configuration. This pattern was similarly observed in SynI-jRCaMP1b recordings: left dCA1 showed a greater pre-entry AUC than right vCA1 in LD↔RV (Figure 4M-N), and right dCA1 showed a greater pre-entry AUC than left vCA1 in RD↔LV (Figure 4R-S), with no significant differences in post-entry AUC, peak amplitude, or peak time (Figure 4O-Q, T-V). In contrast to the cell-type-restricted asymmetry observed during home cage recordings, the dorsal pre-entry bias around coordinated center entries was detected in both pyramidal-cell-biased and pan-neuronal signals, suggesting a behaviorally driven dorsal CA1 recruitment that is broadly distributed across the CA1 network during this behavioral transition. Two implications follow. First, dorsal-ventral CA1 organization is behavior-state-dependent: neither *ex vivo* excitability nor home cage event rate predicts the activity gradient observed around defined exploratory transitions. Second, coordinated center entries preferentially recruit dorsal CA1 before the entry itself, consistent with a contribution of dCA1 to the spatial-contextual evaluation that precedes approach into an exposed area(O’Keefe, 1978; Moser et al., 1995).

**Figure 4.**
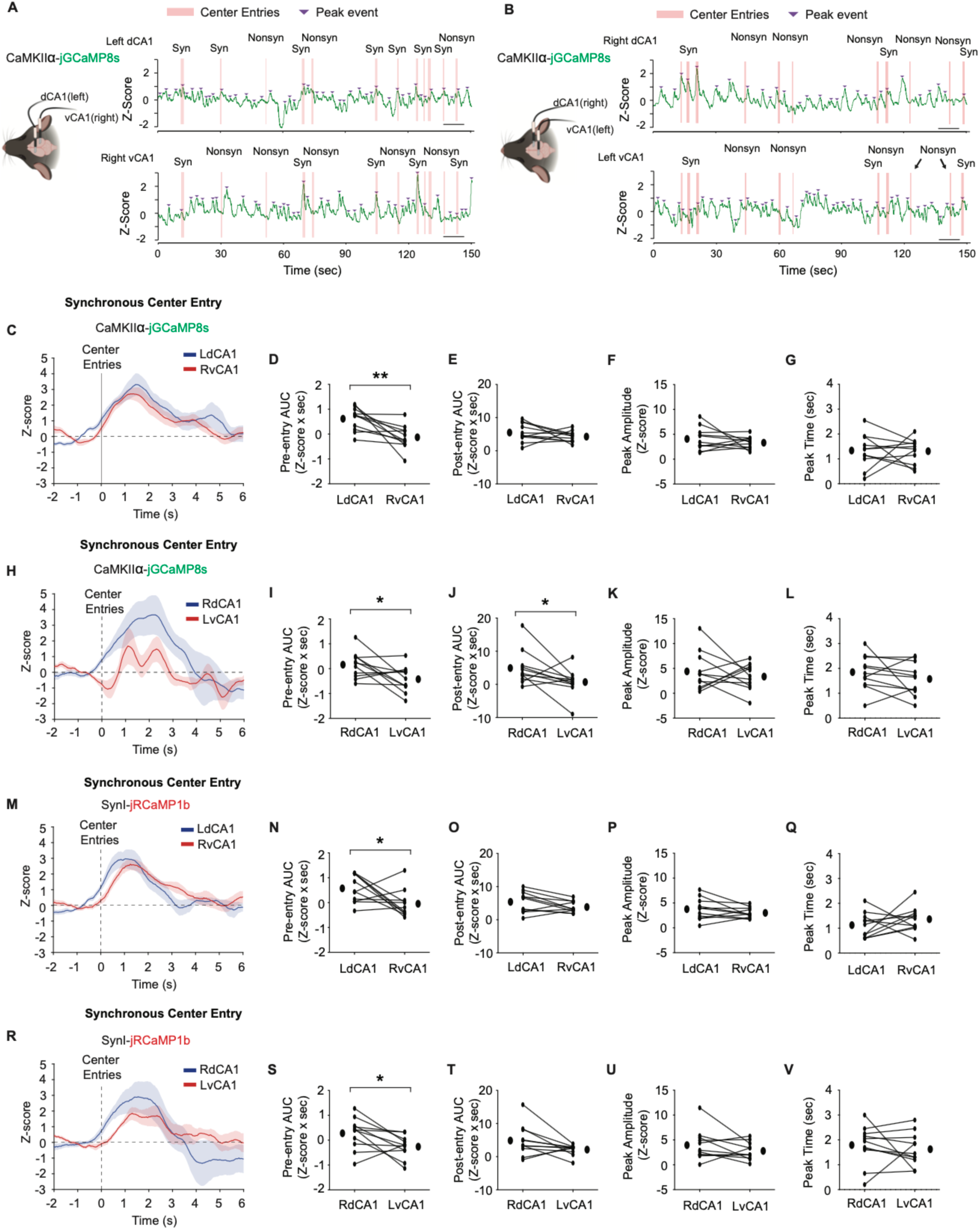
Coordinated open-field center entries recruit dorsal CA1 ahead of ventral CA1. (A, B) Representative CaMKIIα-jGCaMP8s calcium traces during the open-field test from the LdCA1↔RvCA1 (A) and RdCA1↔LvCA1 (B) configurations. Pink bars indicate center entries; purple arrowheads indicate detected calcium events. Center entries were classified as synchronous when peaks were detected in both recorded regions within ± 1 s of entry onset, and as nonsynchronous otherwise; the nonsynchronous subset is analyzed in Figure 5. (C-G) Synchronous center-entry-aligned CaMKIIα-jGCaMP8s responses in the LdCA1↔RvCA1 configuration. Mean Z-scored traces (C) and quantification of pre-entry AUC (D, [-2, 0] s relative to entry), post-entry AUC (E, [0, +6] s), peak amplitude (F), and peak time (G). LdCA1 showed a significantly greater pre-entry AUC than RvCA1; post-entry AUC, peak amplitude, and peak time did not differ. (H-L) Synchronous center-entry-aligned CaMKIIα-jGCaMP8s responses in the RdCA1↔LvCA1 configuration. RdCA1 showed significantly greater pre-entry (I) and post-entry (J) AUC than LvCA1; peak amplitude (K) and peak time (L) did not differ. (M-Q) Synchronous center-entry-aligned SynI-jRCaMP1b responses in the LdCA1↔RvCA1 configuration. LdCA1 showed a significantly greater pre-entry AUC than RvCA1 (N); post-entry AUC (O), peak amplitude (P), and peak time (Q) did not differ. (R-V) Synchronous center-entry-aligned SynI-jRCaMP1b responses in the RdCA1↔LvCA1 configuration. RdCA1 showed a significantly greater pre-entry AUC than LvCA1 (S); post-entry AUC (T), peak amplitude (U), and peak time (V) did not differ. Data are shown as individual mice with mean ± SEM. Lines connect paired dorsal and ventral CA1 measurements from the same animal. Four metrics were tested per configuration (pre-entry AUC, post-entry AUC, peak amplitude, peak time). Pre-entry and post-entry windows were defined as [-1, 0] s and [0, +2] s relative to center entry onset. p-values and test statistics are reported in Table_statistical_analyses. *p < 0.05, **p < 0.01. Sample sizes. LdCA1↔RvCA1, n = 11 mice; RdCA1↔LvCA1, n = 11 mice.

### Nonsynchronous center entries dissociate dorsal and ventral CA1 responses and reveal post-entry ventral suppression

A potential concern with the synchronous-entry analysis is that the dorsal pre-entry bias could reflect locomotion, generic arousal, or center-zone entry per se, rather than a behaviorally meaningful CA1 recruitment pattern. To address this directly, we examined nonsynchronous center entries, entries that meet the same behavioral criterion but are not accompanied by a coordinated calcium event in both recorded regions. If the synchronous-entry response were driven solely by movement into the center, both entry classes should produce similar regional response profiles. They do not. In CaMKIIα-jGCaMP8s recordings, nonsynchronous entries in the LD↔RV configuration produced an attenuated and divergent response (Figure 5A). Left dCA1 retained a significantly greater pre-entry AUC than right vCA1 (Figure 5B), but no significant post-entry difference emerged (Figure 5C). The dorsal pre-entry bias was therefore present in this configuration even when bilateral coordinated recruitment was absent, suggesting that left dCA1 is engaged before center entries that do not propagate to ventral CA1. In the RD↔LV configuration, in contrast, the pre-entry difference between right dCA1 and left vCA1 was abolished (Figure 5D-E), and a significant post-entry difference emerged that was driven by a relative suppression of left vCA1 activity (Figure 5F). This post-entry vCA1 suppression suggests that ventral CA1 does not merely fail to be recruited around uncoordinated center entries but shows a transient negative deflection following entry, a pattern not previously reported in bilateral dorsal-ventral CA1 photometry studies. The same configuration-dependent pattern was observed in SynI-jRCaMP1b recordings: a preserved left-dCA1 pre-entry bias in LD↔RV (Figure 5G–H, with no post-entry difference, Figure 5I), and abolished pre-entry bias but significant post-entry vCA1 suppression in RD↔LV (Figure 5J-L). Because this pattern is present in both indicators, the post-entry vCA1 suppression is unlikely to reflect a CaMKIIα-specific mechanism and may instead reflect a broader vCA1 disengagement signal during exploratory transitions that are not bilaterally coordinated. These results suggest three conclusions. First, the dorsal pre-entry bias around coordinated center entries cannot be reduced to locomotion, center-zone entry, or generalized arousal, because the same behavioral event produces a different regional response profile when bilateral recruitment is absent. Second, ventral CA1 contributes to the open-field response not only by coactivation with dorsal CA1 but also by a transient post-entry suppression in the RD↔LV configuration. This signature warrants analysis in stress-modulated states, where vCA1 disengagement may be expected to change. Third, the persistence of the left-dCA1 pre-entry bias even during nonsynchronous entries is consistent with the home cage finding that left dCA1 is the source of the CaMKIIα-defined hemispheric asymmetry. We note that RvCA1 showed a delayed positive deflection between 4-6 s post-entry in the LD↔RV configuration (Figures 5A, G) that was absent in the RD↔LV pairing; however, because this window likely encompasses exit initiation in addition to center-zone occupancy, and because no analogous pattern was observed for the symmetrical configuration, we do not interpret this late signal within the current analytical framework. Whether this delayed vCA1 response reflects exit-related processing, a configuration-specific vCA1 dynamic, or an artifact of unequal dwell-time distributions across conditions remains to be determined.

**Figure 5.**
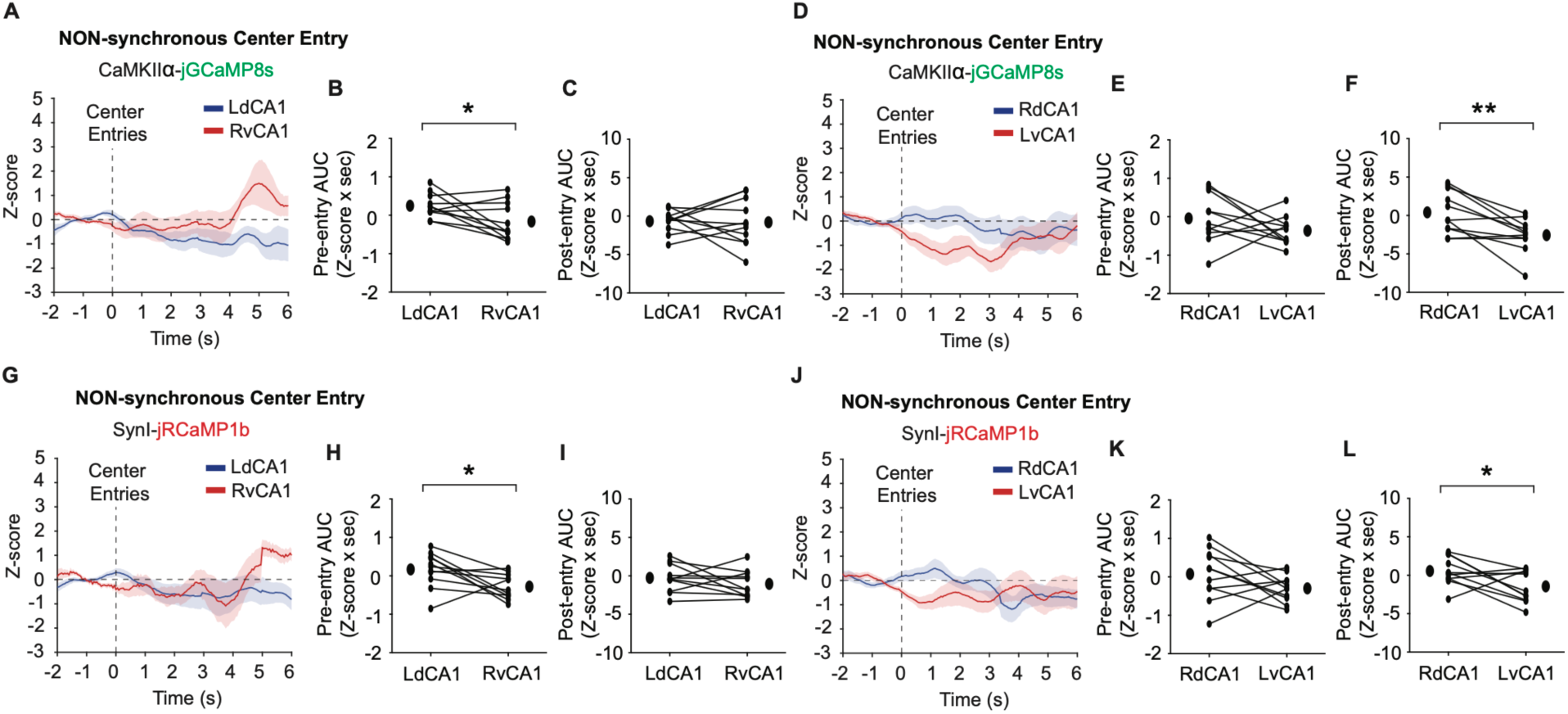
Nonsynchronous center entries dissociate dorsal-ventral CA1 responses and reveal post-entry ventral suppression in the RdCA1↔LvCA1 configuration. (A-C) Nonsynchronous center-entry-aligned CaMKIIα-jGCaMP8s responses in the LdCA1↔RvCA1 configuration. Mean Z-scored traces (A) and quantification of pre-entry AUC (B) and post-entry AUC (C). LdCA1 showed a significantly greater pre-entry AUC than RvCA1; post-entry AUC did not differ. (D-F) Nonsynchronous center-entry-aligned CaMKIIα-jGCaMP8s responses in the RdCA1↔LvCA1 configuration. Mean Z-scored traces (D) and quantification of pre-entry AUC (E) and post-entry AUC (F). Pre-entry AUC did not differ between RdCA1 and LvCA1; post-entry AUC was significantly greater in RdCA1 than in LvCA1, reflecting a relative suppression of LvCA1 activity after center entry. (G-I) Nonsynchronous center-entry-aligned SynI-jRCaMP1b responses in the LdCA1↔RvCA1 configuration. LdCA1 showed a significantly greater pre-entry AUC than RvCA1 (H); post-entry AUC did not differ (I). (J-L) Nonsynchronous center-entry-aligned SynI-jRCaMP1b responses in the RdCA1↔LvCA1 configuration. Pre-entry AUC did not differ (K); post-entry AUC was significantly greater in RdCA1 than in LvCA1 (L). Data are shown as individual mice with mean ± SEM. Lines connect paired dorsal and ventral CA1 measurements from the same animal. Two metrics were tested per configuration (pre-entry AUC, post-entry AUC); Pre-entry and post-entry windows were defined as in Figure 4. p-values and test statistics are reported in Table_statistical_analyses. *p < 0.05, **p < 0.01. Sample sizes. LdCA1↔RvCA1, n = 11 mice; RdCA1↔LvCA1, n = 11 mice; same animals as Figure 4.

## Discussion

The major findings from the present study are that (1) ventral CA1 pyramidal neurons are intrinsically more excitable and synaptically more responsive than dorsal CA1 neurons *ex vivo*; (2) *in vivo*, spontaneous CaMKIIα-defined event rates in dorsal CA1 show a left-biased hemispheric asymmetry not observed in ventral CA1 or in SynI-defined recordings; (3) dorsal-ventral temporal coordination during home cage rest is configuration-dependent and restricted to pyramidal-cell-biased signals; and (4) around open-field center entries, dorsal CA1 is preferentially recruited before entry across both configurations and indicators, whereas non-coordinated entries reveal a post-entry ventral CA1 suppression. Taken together, these results suggest that dorsal-ventral CA1 organization is layered rather than binary, shaped by cell-type identity, hemispheric configuration, and behavioral state, and that a pyramidal-cell-specific left dorsal CA1 asymmetry is a structural feature of this organization.

The *ex vivo* recordings reproduce the core observation that ventral CA1 pyramidal neurons are intrinsically more excitable than their dorsal counterparts, with a more depolarized resting potential, higher input resistance, greater spike output, and steeper AMPAR-mediated EPSP slopes(Dougherty et al., 2012; Kim and Johnston, 2015). HCN-channel-mediated *I*_h_ is a well-documented contributor to this gradient(Dougherty et al., 2013). However, our *in vivo* data show that this cellular gradient does not translate into a uniform ventral-dominant activity pattern under freely behaving conditions. If intrinsic excitability were the dominant determinant of *in vivo* event rates, ventral CA1 should have shown elevated rates in both contralateral configurations and in both indicators. It did not. This dissociation suggests that *ex vivo* excitability is best understood as a cellular substrate rather than a direct prediction of *in vivo* population activity, and it is consistent with our prior work showing that dorsal CA1 carries its own stress-sensitive plasticity, including HCN1/*I*_h_ changes that contribute to behavioral susceptibility after social defeat stress(Kim et al., 2022; Kim et al., 2025). Baseline dorsal-ventral excitability and stress-induced dorsal CA1 plasticity may therefore represent distinct but interacting levels of hippocampal specialization.

A central finding of this study is that the dorsal-ventral organization of *in vivo* CA1 activity depended on which neuronal population was sampled. CaMKIIα-jGCaMP8s recordings revealed configuration-dependent differences in spontaneous event rate and inter-regional synchrony, whereas SynI-jRCaMP1b recordings did not. Because the SynI promoter drives expression in both excitatory and inhibitory populations(Nathanson et al., 2009), the SynI signal averages over heterogeneous cell types whose individual activities can move in opposite directions during the same behavioral epoch(Klausberger and Somogyi, 2008). Population-averaged signals can therefore mask selective changes in pyramidal ensembles even when those changes are reproducible. This may also help explain why some *in vivo* studies have not found straightforward dorsal-ventral differences in mean firing rate despite robust *ex vivo* excitability gradients(Jung et al., 1994; Chockanathan and Padmanabhan, 2021). Consistent with this interpretation, supplementary analyses of peak amplitude and median absolute deviation did not reveal consistent regional or hemispheric effects in either indicator, suggesting that the major organizational differences were carried by event occurrence rather than by event size or baseline signal variability. The hemispheric asymmetry we observed is not, in our view, attributable to recording artifacts. Left dorsal CA1 showed higher CaMKIIα-defined event number and frequency than right dorsal CA1, whereas ventral CA1 did not show an analogous left-right difference, and the strongest dorsal-ventral separation appeared in the RD↔LV configuration. These observations are consistent with a now-substantial literature showing that hippocampal function is not symmetric across hemispheres. CA3-to-CA1 synapses originating from the left and right hemispheres differ in synaptic morphology, glutamate receptor composition, and plasticity(Shinohara et al., 2008; Kohl et al., 2011; Shipton et al., 2014), and behavioral studies have shown that long-term spatial memory depends preferentially on left CA3–CA1 synaptic transmission(Shipton et al., 2014). Our data extend this concept by suggesting that hemispheric context shapes how dorsal-ventral CA1 dynamics are expressed *in vivo*. This interpretation should remain cautious, however, because left-right comparisons in the present study were inferred across contralateral recording configurations rather than from simultaneous bilateral recordings of the same dorsal or ventral subregion. Direct simultaneous LdCA1–RdCA1 and LvCA1–RvCA1 recordings will be needed to distinguish a true hemispheric specialization from configuration-specific effects.

The lead–lag analysis indicates that dorsal-ventral organization is not defined only by event rate but also by how events are temporally coordinated. In the CaMKIIα signal, the RD↔LV configuration showed reduced synchronous events and increased ventral CA1 solo events, whereas dorsal-leading and ventral-leading proportions were unchanged. The dominant temporal effect was therefore a loss of bilateral coupling and a gain of independent ventral CA1 activity, rather than a directional shift in temporal precedence. This effect was absent in SynI recordings. Because fiber photometry has slower kinetics than spike-resolved electrophysiology, these classifications describe temporal precedence at the population level and not synaptic or causal directionality(Ali and Kwan, 2020; Simpson et al., 2024). With this caveat in mind, the reproducibility of the synchrony and solo-event differences suggests that dorsal and ventral CA1 are not independently active regions but components of a coordinated longitudinal network whose coupling is shaped by hemisphere and cell type.

The open-field data show that this organization reorganizes around behaviorally defined transitions. During synchronous center entries, dorsal CA1 showed a greater pre-entry response than ventral CA1 in both configurations and indicators, suggesting a dCA1 contribution to the spatial-contextual evaluation that precedes approach into an exposed area. This finding is consistent with Jimenez et al. (2018), who identified ventral CA1 anxiety cells preferentially active during occupancy of anxiogenic environments. Two features of that study warrant direct comparison. First, Jimenez et al. performed the open field under bright illumination (∼650 lux), emphasizing the anxiogenic value of the center; our recordings were conducted under low illumination (6.6 lux), under which center-zone entry is a transition point during exploration rather than a strong anxiogenic load. Second, Jimenez et al. reported that vCA1 anxiety cells increased activity largely after entry, not before. Our findings are therefore not in conflict with theirs: the pre-entry dorsal CA1 response and the post-entry vCA1 response of Jimenez et al. reflect the transition phase and occupancy phase of the same behavioral event, respectively, and are complementary rather than competing. Dorsal and ventral hippocampus are increasingly understood to operate along overlapping functional gradients rather than as fully separable spatial-versus-emotional modules (Fanselow and Dong, 2010; Strange et al., 2014), and our pre-entry dorsal CA1 recruitment is consistent with that framework. The nonsynchronous center-entry analysis provides an important internal control for these findings. Because synchronous and nonsynchronous entries share the same behavioral definition, the finding that they produce different regional response profiles suggests that the synchronous-entry response reflects coordinated circuit engagement rather than a movement- or arousal-driven artifact. Interestingly, nonsynchronous entries in the RD↔LV configuration revealed a post-entry suppression of left vCA1, a pattern present in both CaMKIIα and SynI recordings and therefore unlikely to reflect a CaMKIIα-specific mechanism. This post-entry vCA1 suppression may reflect a broader vCA1 disengagement signal during uncoordinated exploratory transitions. The persistence of the left-dCA1 pre-entry bias even during nonsynchronous entries is consistent with the home cage finding that left dCA1 is the source of the CaMKIIα-defined hemispheric asymmetry: left dCA1 is recruited before center entries regardless of whether that recruitment propagates to ventral CA1.

Several limitations should be acknowledged. First, CaMKIIα- and SynI-driven indicators provide separation between pyramidal-cell-biased and broadly neuronal signals but do not resolve interneuron subclasses, projection-defined pyramidal populations, or non-neuronal contributions. Second, fiber photometry captures population-level fluorescence and cannot distinguish the recruitment of additional neurons from changes in per-neuron firing rate or per-spike calcium influx. Third, lead–lag analysis from photometry data describes temporal precedence at the population level but cannot establish synaptic directionality. Fourth, a scope-limit of the classifier itself should be noted: the algorithm matches positive-going events between channels and captures co-activation but not anti-phase coupling. A D-up/V-down or D-down/V-up event is categorized as a solo or nonsynchronous event rather than as a distinct anti-correlated class. This may be relevant to the post-entry vCA1 suppression in the RD↔LV configuration (Figure 5F), which is consistent with an anti-phase pattern that the current classifier does not explicitly resolve. Fifth, hemispheric asymmetry here was inferred from contralateral recording configurations rather than simultaneous bilateral recordings of the same subregion.

In summary, dorsal-ventral CA1 organization *in vivo* is a layered, state-dependent phenomenon shaped by hemispheric configuration and neuronal population identity. A CaMKIIα-specific left dorsal CA1 asymmetry, absent in ventral CA1 and in SynI-defined signals, accounts for apparent dorsal-ventral differences during home cage rest and shapes inter-regional temporal coupling. During coordinated center entries, dorsal CA1 is preferentially recruited before the behavioral transition, whereas uncoordinated entries reveal a post-entry vCA1 suppression. These results establish a baseline framework for asking how this organization is reshaped under chronic stress, where hemisphere-specific dorsal CA1 plasticity and vCA1 disengagement during exploratory transitions are likely to be the most informative dimensions to examine.

## Acknowledgements

We thank Dr. Jiwon Kim for assistance with virus injection. This work was supported by a start-up fund (to CSK) from the Medical College of Georgia at Augusta University, and by NIH grant R01MH134958 (to CSK) and R01MH137204 (to SK).

## Author Contributions

**C.S.K.** conceived the project, designed the experiments, performed *ex vivo* electrophysiology and *in vivo* fiber photometry recordings, analyzed all data, and wrote the manuscript. **J.B.** performed *in vivo* fiber photometry experiments. **M.L.** performed behavioral experiments. **S.K.** provided technical guidance during the early stage of the *in vivo* fiber photometry experiments. All authors reviewed and edited the manuscript.

## Data availability

The data that support the findings of this study are available from the corresponding author upon reasonable request.

