## Supplemental information for "Pyramidal-cell-specific hemispheric asymmetry shapes dorsoventral CA1 dynamics during rest and exploratory behavior"

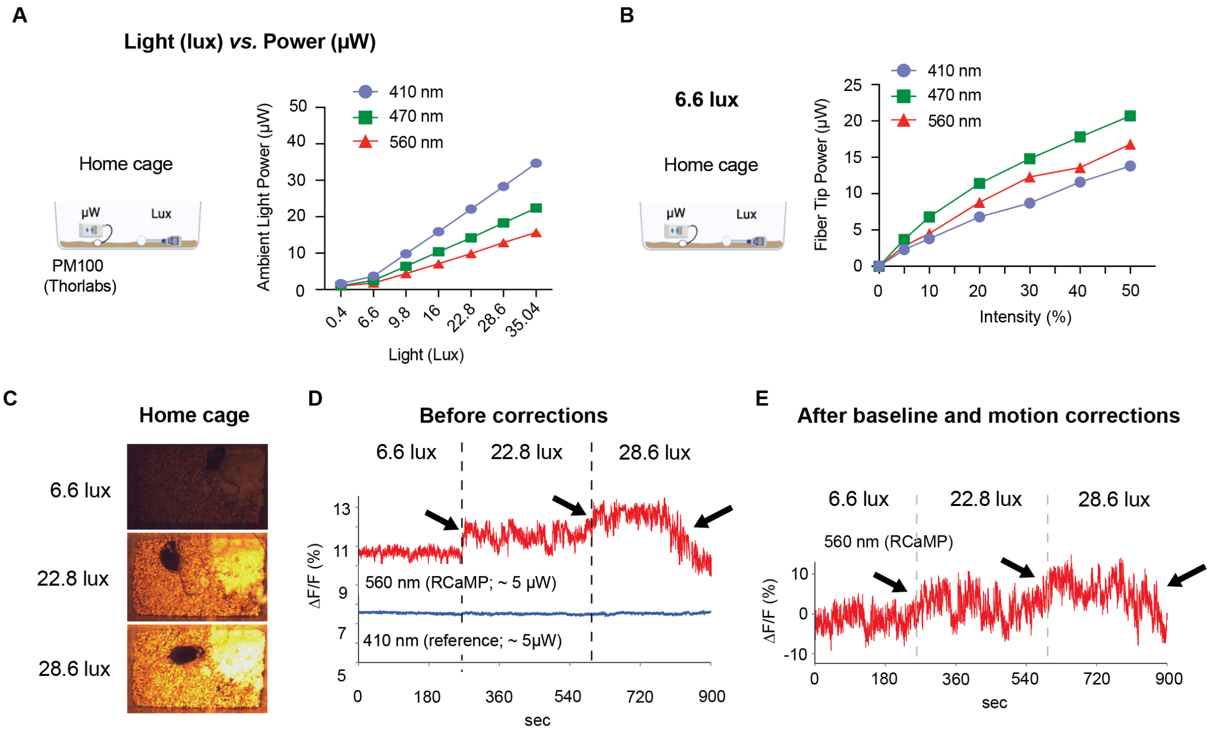


**Supplementary Figure S1. Effects of ambient light intensity on fiber photometry signal stability and fluorescence measurements.** (A) Relationship between ambient light intensity (lux) and measured optical power (μW) across excitation wavelengths (410 nm, 470 nm, and 560 nm). Increasing environmental illumination resulted in wavelength-dependent increases in detected optical power, indicating potential light contamination of recorded signals. (B) Output power calibration curves for individual photometry channels (CH1 and CH2) at low ambient light conditions (6.6 lux), showing stable and comparable light delivery across excitation wavelengths over a range of stimulation intensities. (C) Representative images of home cage conditions under different ambient light levels (6.6, 22.8, and 28.6 lux), illustrating increased environmental brightness and potential for reflected light contamination at higher illumination levels. (D) Representative raw fluorescence traces (ΔF/F) prior to baseline and motion correction under increasing ambient light conditions. Elevated illumination produced baseline shifts and signal drift in the 560 nm (RCaMP) channel, while the 410 nm reference signal remained relatively stable. (E) Fluorescence traces after baseline and motion correction. Although correction procedures partially reduced light-induced artifacts, residual fluctuations and distortions persisted at higher light levels (arrows), indicating incomplete removal of illumination-dependent effects. These results demonstrate that ambient light significantly influences fiber photometry signal stability and highlight the importance of performing recordings under low-light conditions to minimize optical artifacts.

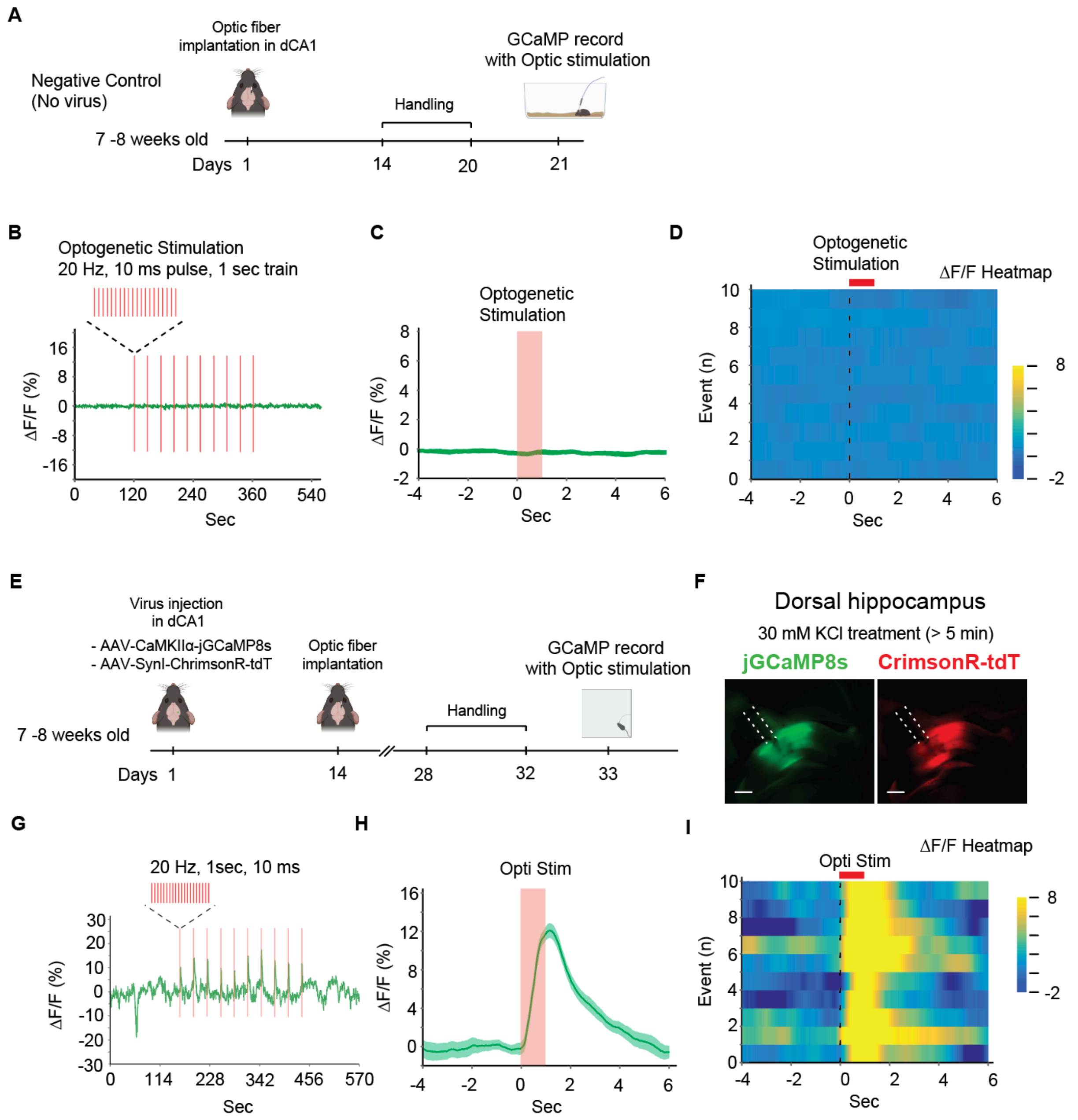


**Supplementary Figure S2. Optogenetic validation of GCaMP signals and empirical basis for calcium event detection parameters.** (A-D) Negative control (no virus). (A) Experimental timeline for negative control animals receiving optic fiber implantation without viral injection. (B) Representative ΔF/F trace during optogenetic stimulation (20 Hz, 10 ms pulse width, 1 s train) showing no stimulus-locked fluorescence response. (C) Event-aligned average trace (mean ± SEM) demonstrating absence of calcium signal change during stimulation. (D) Event-aligned heatmap showing no consistent stimulus-locked fluorescence across trials. (E-I) Optogenetic validation of calcium signals. (E) Experimental timeline for optogenetic validation cohort receiving AAV-CaMKIIα-jGCaMP8s and AAV-Syn-ChrimsonR-tdTomato injections in dCA1 followed by optic fiber implantation. (F) Representative *ex vivo* validation following 30 mM KCl application (>5 min), showing robust fluorescence responses in jGCaMP8s (green) and ChrimsonR-tdTomato (red), confirming functional indicator expression and targeting. (G) Representative ΔF/F trace showing reliable stimulus-locked calcium responses during optogenetic stimulation (20 Hz, 10 ms pulse width, 1 s train). (H) Event-aligned average trace (mean ± SEM) demonstrating a rapid rise and prolonged decay of the evoked calcium transient. (I) Event-aligned heatmap illustrating consistent stimulus-locked fluorescence increases across trials.

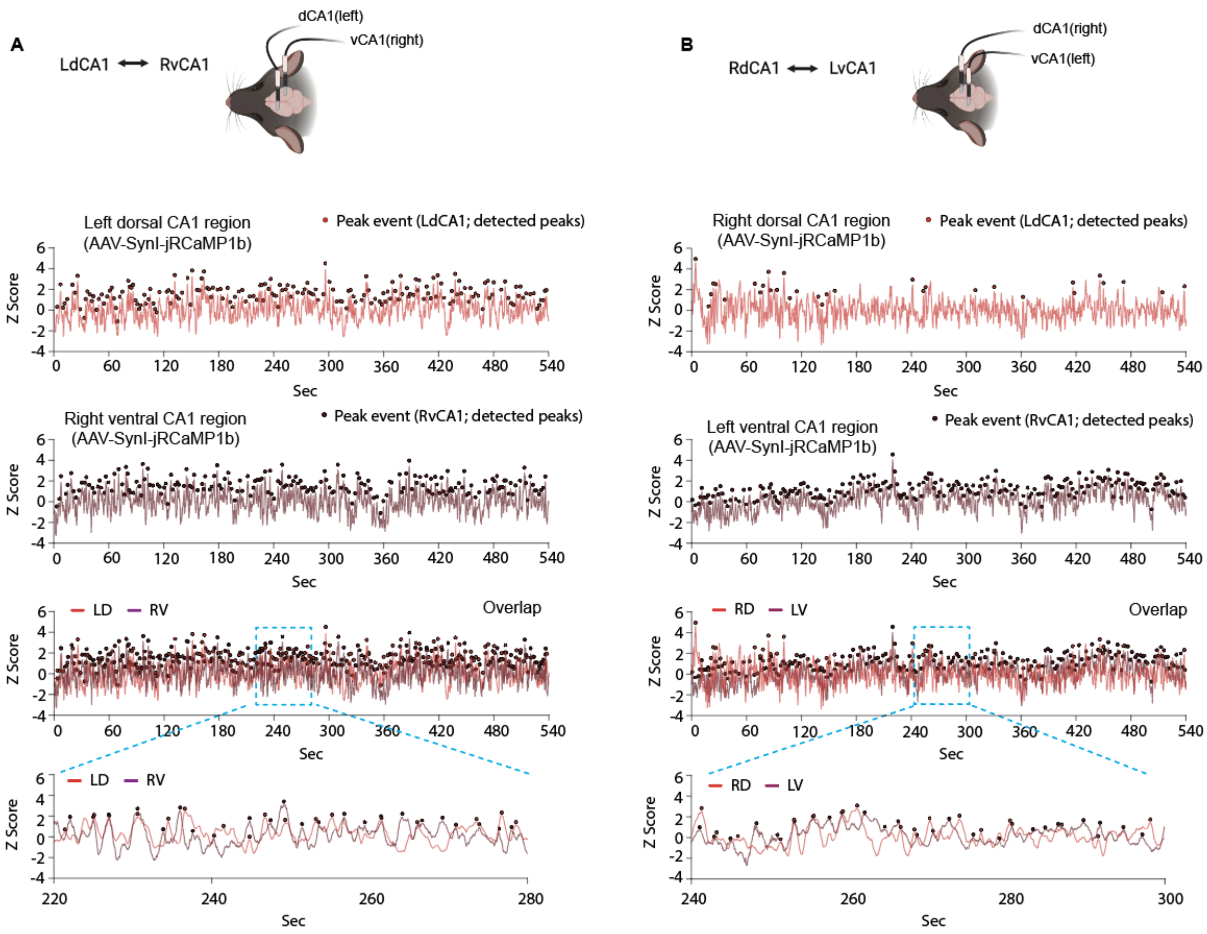


**Supplementary Figure S3. Representative pan-neuronal (Syn1-driven) calcium activity in dorsal and ventral CA1 under home cage conditions.** (A) Representative calcium traces from the LD–RV configuration showing simultaneous recordings of left dorsal CA1 (LdCA1) and right ventral CA1 (RvCA1) using AAV-Syn1-jRCaMP1b. Detected peak events are indicated for each region. Overlaid traces illustrate temporal correspondence between dorsal and ventral signals, and the dashed box highlights a representative time window shown at higher temporal resolution below. (B) Representative calcium traces from the RD–LV configuration showing simultaneous recordings of right dorsal CA1 (RdCA1) and left ventral CA1 (LvCA1). As in (A), detected peak events are indicated, and overlaid traces demonstrate partially overlapping but non-identical activity patterns between regions. Expanded views (dashed box) illustrate fine-scale temporal relationships between dorsal and ventral CA1 signals. Across both configurations, Syn1-driven signals exhibit broadly similar activity patterns between dorsal and ventral CA1, with substantial temporal overlap and no consistent regional bias in event occurrence, consistent with the absence of dorsal–ventral differences observed in quantitative analyses (Figure 2).

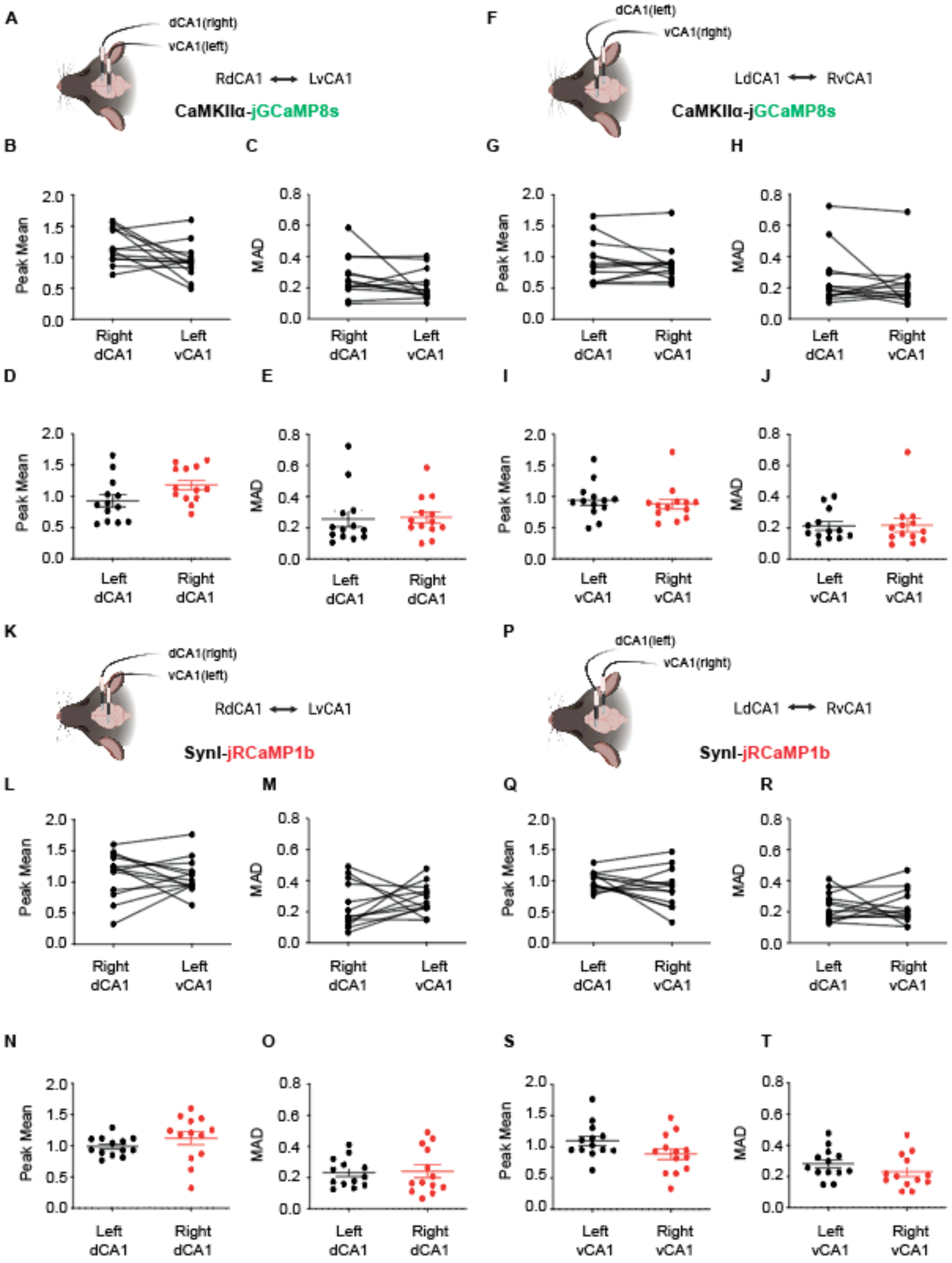


**Supplementary Figure S4. Peak amplitude and signal variability of spontaneous calcium events across contralateral recording configurations.** Peak mean and median absolute deviation (MAD) of detected calcium events during home cage behavior, complementary to the event-rate analyses shown in Figure 2. (A-E) CaMKIIα-jGCaMP8s, RdCA1↔LvCA1 configuration: schematic (A), within-animal paired comparison of right dCA1 vs. left vCA1 peak mean (B) and MAD (C), and inter-animal hemispheric comparison of left vs. right dCA1 peak mean (D) and MAD (E). (F–J) CaMKIIα-jGCaMP8s, LdCA1↔RvCA1 configuration: schematic (F), within-animal paired comparison of left dCA1 vs. right vCA1 peak mean (G) and MAD (H), and inter-animal hemispheric comparison of left vs. right vCA1 peak mean (I) and MAD (J). (K-O) SynI-jRCaMP1b, RdCA1↔LvCA1 configuration: schematic (K), paired peak mean (L) and MAD (M), and hemispheric peak mean (N) and MAD (O). (P-T) SynI-jRCaMP1b, LdCA1↔RvCA1 configuration: schematic (P), paired peak mean (Q) and MAD (R), and hemispheric peak mean (S) and MAD (T). Across both indicators and configurations, peak mean and MAD did not show consistent dorsal-ventral or hemispheric differences, indicating that the regional and configuration-dependent effects observed in home cage recordings (Figure 2) were primarily reflected in event number and frequency rather than peak amplitude or signal variability. Data points represent individual animals; connected lines indicate paired dorsal-ventral measurements from the same animal. Mean ± SEM shown. Paired comparisons: two-tailed paired t-tests. Inter-animal hemispheric comparisons: two-tailed Welch's t-tests. n = 13 mice per configuration.

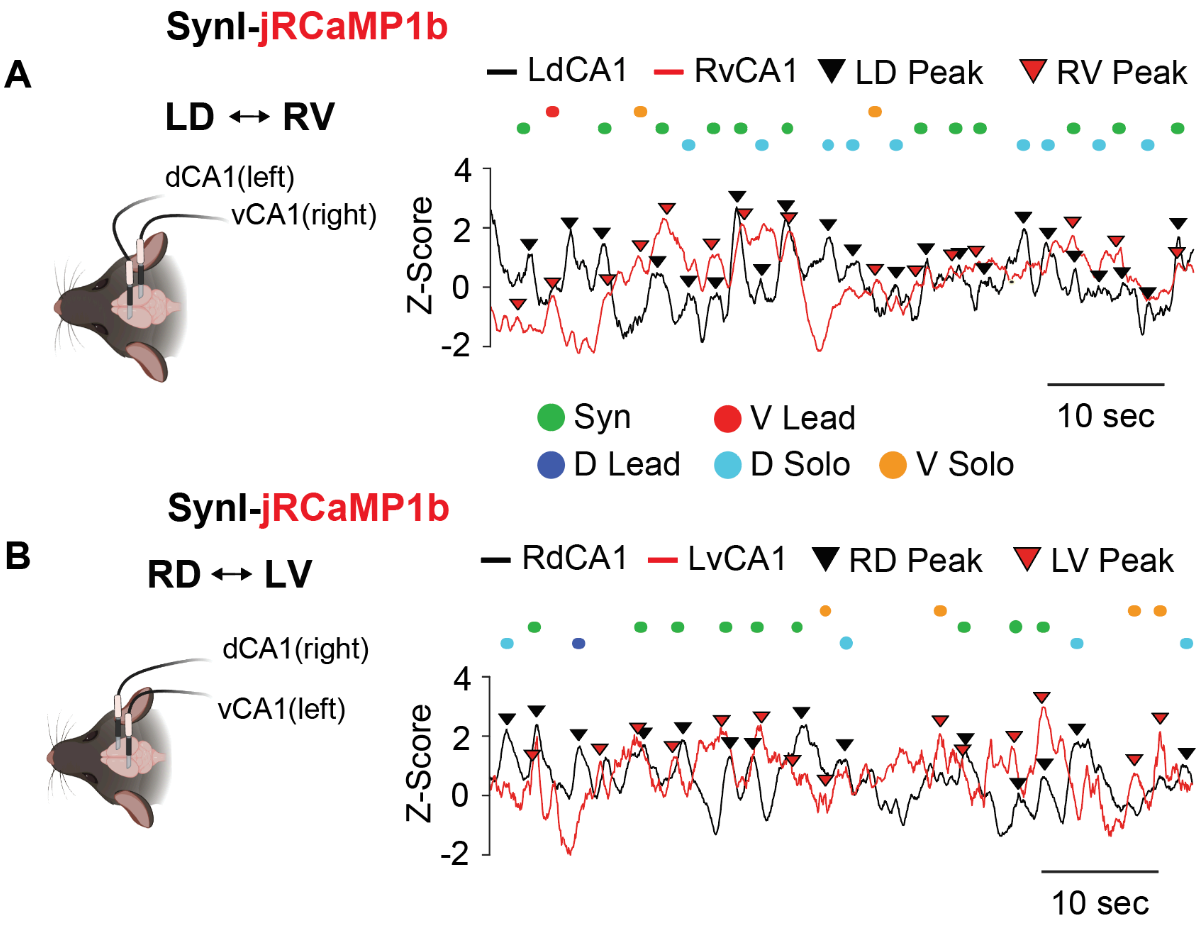


**Supplementary Figure S5. Representative SynI-jRCaMP1b traces with classified calcium events during home cage recordings.** Representative 60-s Z-scored SynI-jRCaMP1b calcium traces recorded simultaneously from dorsal and ventral CA1 during home cage behavior, complementary to the CaMKIIα-jGCaMP8s traces shown in Figure 3A. (A) LD↔RV configuration: simultaneous recordings from left dorsal CA1 (LdCA1, black) and right ventral CA1 (RvCA1, red). (B) RD↔LV configuration: simultaneous recordings from right dorsal CA1 (RdCA1, black) and left ventral CA1 (LvCA1, red). Black and red downward triangles indicate detected calcium events in the dorsal and ventral channels, respectively. Colored dots above each trace indicate the classification of each detected event using the reciprocal nearest-event matching algorithm described in the Methods: synchronous (Syn, green; $|\Delta t| \leq 1$ s), dorsal-leading (D Lead, dark blue; vCA1 follows dCA1 by 1–2 s), ventral-leading (V Lead, red; dCA1 follows vCA1 by 1–2 s), dorsal CA1 solo (D Solo, light blue; no reciprocal partner within ±2 s), and ventral CA1 solo (V Solo, orange; no reciprocal partner within ±2 s). Group-level quantification of these event categories is shown in Figure 3B (right panels) and Figure 3C–H.

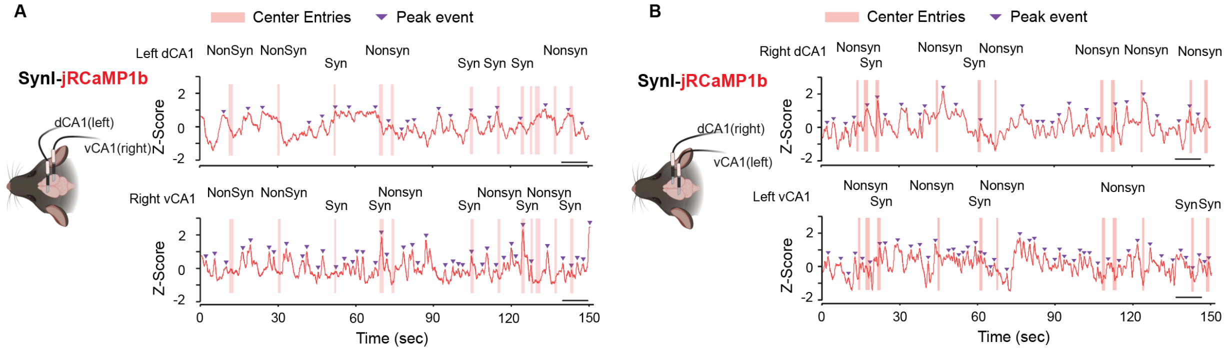


**Supplementary Figure S6. Representative SynI-jRCaMP1b traces during open-field exploration with synchronous and nonsynchronous center entries.** Representative SynI-jRCaMP1b calcium traces from dorsal and ventral CA1 recorded simultaneously during open-field exploration, complementary to the CaMKIIα-jGCaMP8s traces shown in Figure 4A-B. (A) LD↔RV configuration: simultaneous recordings from left dorsal CA1 (LdCA1, top) and right ventral CA1 (RvCA1, bottom). Scare bar: 10 sec. (B) RD↔LV configuration: simultaneous recordings from right dorsal CA1 (RdCA1, top) and left ventral CA1 (LvCA1, bottom). Scare bar: 10 sec. Pink shaded bars mark center-zone entries, and purple downward triangles mark detected calcium events. Each center entry is labeled above the trace as either synchronous (Syn; peaks detected in both recorded regions within ±1 s of entry onset) or nonsynchronous (Nonsyn; absence of bilateral coordination within the same window). Event-aligned quantification of synchronous and nonsynchronous center entries is shown in Figure 4M-V and Figure 5G-L, respectively.
